# Flax latitudinal adaptation at *LuTFL1* altered architecture and promoted fiber production

**DOI:** 10.1101/178772

**Authors:** Rafal M Gutaker, Maricris Zaidem, Yong-Bi Fu, Axel Diederichsen, Oliver Smith, Roselyn Ware, Robin G Allaby

## Abstract

After domestication in the Near East around 10,000 years ago several founder crops, flax included, spread to European latitudes. On reaching northerly latitudes the architecture of domesticated flax became more suitable to fiber production over oil, with longer stems, smaller seeds and fewer axillary branches. Latitudinal adaptations in crops typically result in changes in flowering time, often involving the PEBP family of genes that also have the potential to influence plant architecture. Two PEBP family genes in the flax genome, *LuTFL1* and *LuTFL2*, vary in wild and cultivated flax over latitudinal range with cultivated flax receiving *LuTFL1* alleles from northerly wild flax populations. Compared to a background of population structure of flaxes over latitude, the *LuTFL1* alleles display a level of differentiation that is consistent with selection for an allele III in the north. We demonstrate through heterologous expression in *Arabidopsis thaliana* that *LuTFL1* is a functional homolog of *TFL1* in *A. thaliana* capable of changing both flowering time and plant architecture. We conclude that specialized fiber flax types could have formed as a consequence of a natural adaptation of cultivated flax to higher latitudes.

---

Crop plants were domesticated in the late Pleistocene to early Holocene at multiple centers around the world and many of them subsequently expanded northwards after glacial retreat^1^. The southwest Asian ‘Neolithic Package’^2^ including wheat, barley, lentils, chickpeas and flax was domesticated in the Near East and spread out into Europe^3^. The environmental conditions associated with the higher latitudes of Europe presented the plants of Near Eastern origin with major challenges that have been associated with repeated agricultural collapse^4,5^, possibly due to the limited rates at which adaptation could take place due to the associated substitution load^6^. Plants of Neolithic Package, adapted to Near Eastern conditions, are often long-day and vernalization dependent for flowering. Those crops would have struggled in short growing season in Europe. Typically, wheat and barley adapted to these conditions through photoperiod insensitivity which affects their flowering time^7–12^. Some northern flax types also show similar loss of photoperiodical sensitivity phenotypes^13^. However, in contrast to the Neolithic cereals, flax had a potential advantage in that it could have assimilated adaptations from its wild progenitor species, which extends in range to northwestern Europe and shows patterns of extensive local adaptation^14,15^.

Cultivated flax (*Linum usitatissimum* L.) is a predominantly self-pollinating^16^, diploid crop plant that was domesticated from pale flax (*L. bienne* Mill.)^17^ in the Near East^2^. Originally domesticated for its oily seeds^18^, flax is currently grown as either an oil or fiber crop. Specialized fiber varieties likely emerged during the Neolithic ‘Textile Revolution’^19–21^, which started in Central Europe about 6,000 years Before Present (BP). The two flax forms are different in their seed size and distinct plant architecture^22^. In particular, specialized fiber varieties are characterized with lower seed weight, fewer axillary branches and higher technical stem length, relative to oil varieties^23,24^. It has been shown that those traits are under divergent selection in both varieties^25^.

In *Arabidopsis thaliana*, plant architecture can be modified simultaneously with both flowering time by changes in regulation of the *FT* and *TFL1* genes^26^, which belong to the phosphatidyl ethanolamine-binding protein (PEBP) family^27^ and control plant development through regulation of floral meristem initiation. An allelic variant of a PEBP gene in barley, *HvCET* has been involved in adapting this crop to the European climate through later flowering, suggesting the potential importance of this family in latitudinal adaptations of crops during the spread of agriculture^10^. Furthermore, *FT* and *TFL1* can control plant stature through promotion or delaying floral initiation^26^ and are highly conserved in eudicots^28^. The orthologs of *FT* and *TFL1* in tomato have a strong effect on tomato fruit weight and number^29^, plant architecture and height, and flowering time^30,31^. Such differences in fruit weight and plant stature underlie the distinction between fiber and oil specialized varieties in flax, raising the possibility that the PEBP gene family in flax could be associated with both architectural changes and adaptation to northerly latitudes through modifying flowering time.

In this study we screened PEBP gene orthologs in flax for signatures of selection along increasing latitudes in European wild and cultivated flax. We additionally genotyped restriction site-associated regions of flax genome to understand the underlying population structure in flax. Finally, we validated the effect of a gene candidate for selection, *LuFTL1* on plant architecture and flowering time through heterologous expression in *Arabidopsis thaliana*.

## Results

### PEBP orthologs in flax and the diversity at *LuTFL1* locus

In order to investigate the role of PEBP genes in latitudinal adaptation and architecture control in cultivated flax, we surveyed orthologs of this gene family. A scan for homologs of *A. thaliana* PEBP genes in the flax assembly^32^ revealed eight loci of interest with architectures similar to *TFL1* gene (Supplementary Figure S1; Supplementary Table S1). Out of those, seven loci were amplified and sequenced from flax genomic DNA (Supplementary Figure S2), four could be easily re-sequenced without cloning and showed conserved four-exon structure (Supplementary Figure S1). Phylogenetic analyses comparing the flax, *A. thaliana* and Lombardy poplar orthologs revealed that three putative PEBP members in flax are closely related to floral suppressors TFL1 and ATC, while one has similarity to their antagonist, FT (Supplementary Figure S3). Resequencing of those genes in six accessions of flax indicated high levels of polymorphism in pale flax at the *LuTFL1* locus, suggestive of broad standing variation.

We re-sequenced the *LuTFL1* locus in 32 wild and 113 cultivated flax accessions (Supplementary Figure S4A, Supplementary Table S2). For comparison, we also re-sequenced the *LuTFL2* locus in the same population of flaxes. We discovered wide molecular diversity at both loci, with 16 and 11 alleles in *LuTFL1* and *LuTFL2* respectively (Supplementary Table S3). A phylogenetic network of *LuTFL1* shows that allelic diversity is greater in pale flax (top plane, Figure 1A) when compared to cultivated flax (bottom plane, Figure 1A). The majority (100 out of 103) of cultivated flaxes were associated with just two of the groups (I and III) also found in the wild. There were several groups apomorphic to groups I and III that appeared only in our cultivated samples that likely represent post-domestication diversification (alleles XI, XII, XIII, XIV, XV and XVI). Interestingly, the *LuTFL2* network shows greater diversity in cultivated flax allele compared to pale flax (Supplementary Figure S5A, Supplementary Table S3).

**Figure 1.**
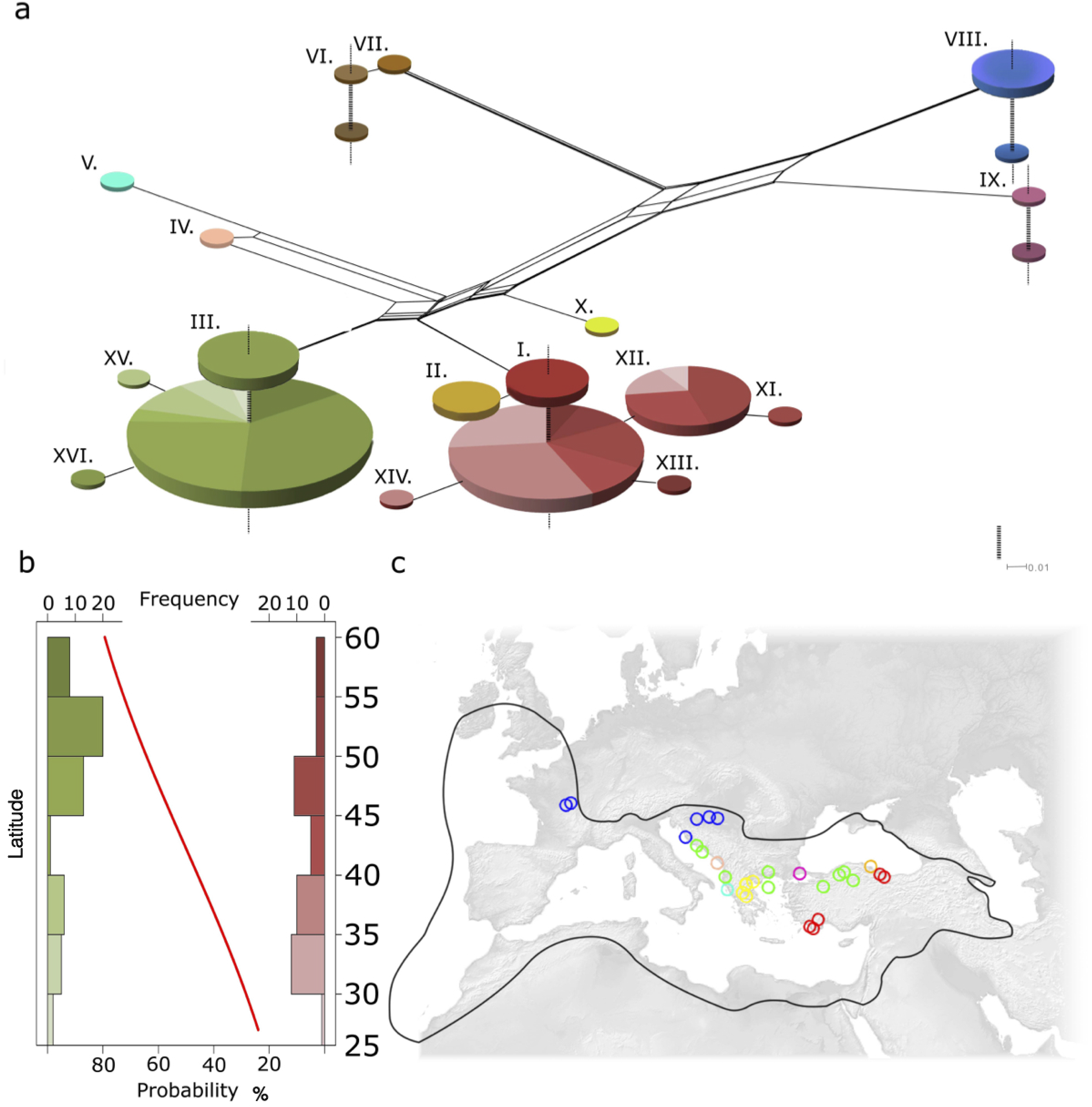
Haplotype network of *LuTFL1* locus in cultivated and pale flax: A. Splits Tree network of pale (top) and cultivated (bottom) flax, size of nodes is proportional to number of samples with the same haplotype, continuous branches denote molecular distance between haplotypes, vertical dotted lines link different species within same haplotype. B. Histogram showing latitudinal gradient of *LuTFL1* alleles in cultivated flax, in green, frequency of northern haplotype cluster (*LuTFL1*.III), in red, frequency of southern haplotype cluster (*LuTFL1*.I) with fitted logistic regression curve (p-value of 0.00144) reflecting occurrence probability of northern haplotype in latitude gradient. C. Map of Europe marked with wild distribution of pale flax (black line) and pale flax sampling locations (colours correspond to haplotypes in splits network).

There are multiple alleles in both *LuTFL1* and *LuTFL2* that are shared by pale and cultivated flaxes. Such a pattern could be a result of multiple alleles being included during domestication or through post-domestication gene flow from wild to domestic species. The phylogeographic distribution of *LuTFL1* alleles in pale flax (Figure 1C), supports the notion that allele I was passed from pale flax to cultivated flax during the initial domestication process in the Near East. Conversely, in pale flax allele III is associated with the Bosphorus strait and the Balkans beyond the domestication area but not in Eastern or Southern Turkey. We conclude that *LuTFL1* III likely entered the cultivated flax gene pool through gene flow from pale flax during the agricultural expansion into Europe, and reached higher frequencies in more northerly latitudes (Figure 1B). The high diversity of *LuTFL2* alleles present in the cultivated gene pool but absent from the wild gene pool suggests that they may have originated from wild populations outside our sample set, or have become extinct in the wild. Despite not being closely located in the genome, allele X of *LuTFL2* occurs in a strong disequilibrium with allele III of *LuTFL1* (Supplementary Table S4), shows a similar latitudinal gradient (Supplementary Figure S5B), and is absent from cultivated flaxes carrying the *LuTFL1* allele I.

The significant latitudinal clines of *LuTFL1* alleles I more frequent in the south and III more frequent in the north (Figure 1B) are more evident in the historic landraces (Supplementary Figure S6). A notable exception to this trend is allele XII from the group I associated alleles which occurs at higher frequencies at higher latitudes, but is absent from the historic samples. We considered populations as ‘northern’ or ‘southern’ relative to 40th circle of latitude of northern hemisphere for calculation of fixation index statistics F_ST_=0.26, which indicated high population differentiation at *LuTFL1* locus. Enrichment of *LuTFL1* III in the north could be a result of drift combined with isolation by distance (IBD), or alternatively it could be an effect of selection acting on adaptive alleles.

We tested if *LuTFL1* and *LuTFL2* alleles are distributed in the frequencies that match the expectation under neutral evolution using Tajima’s D, R2, Fu and Li’s D_2_ and F methods (Supplementary Table S5). Based on *LuTFL2* data we could not reject the neutrality hypothesis, however, significantly negative scores in *LuTFL1* tests indicated an excess of rare alleles in cultivated flax. Such a pattern of molecular diversity could be explained either by population structure, demographic effects such as expansion, or selection. To distinguish between these possibilities, we investigated the population structure and allele frequencies at neutral loci through a restriction site-associated (RAD) sequencing approach^33,34^.

### Population structure from genome-wide SNPs in wild and pale flax

We surveyed the genomes of 90 accessions of pale (28) and cultivated (62) flax (Supplementary Table S2) by RAD sequencing^33^. We used *de novo* assembly approach on sequenced fragments^35^, from which we found 993 polymorphic RAD tags encompassing the total of 1686 SNPs present in at least 80% accessions. Based on those, we calculated genetic distances, which were used for multidimensional scaling analysis. Three main clusters that correspond to three populations were revealed: cultivated flax, dehiscent flax and pale flax (Figure 2A). Using an ADMIXTURE^36^ approach we identified four distinct ancestral components (lowest 5-fold cross validation error for K=4). The pale flax of Eastern Anatolia represents all the admixture components that are present in cultivated flaxes, including dehiscent varieties (Figure 2B), which supports an eastern population involvement with the original domestication process. Two pale flaxes located in eastern Greece outside the domestication center show sizeable admixture with dehiscent varieties. This is congruent with previous reports that identify dehiscent varieties as genetically distinct from other domesticated flax varieties, possibly reflecting an earlier independent domestication^37,38^. A latitudinal gradient is apparent in the cultivated flax types, evident from a tendency for southern populations to be affiliated to an orange grouping, and a cyan grouping in northern populations in Figure 2B.

**Figure 2.**
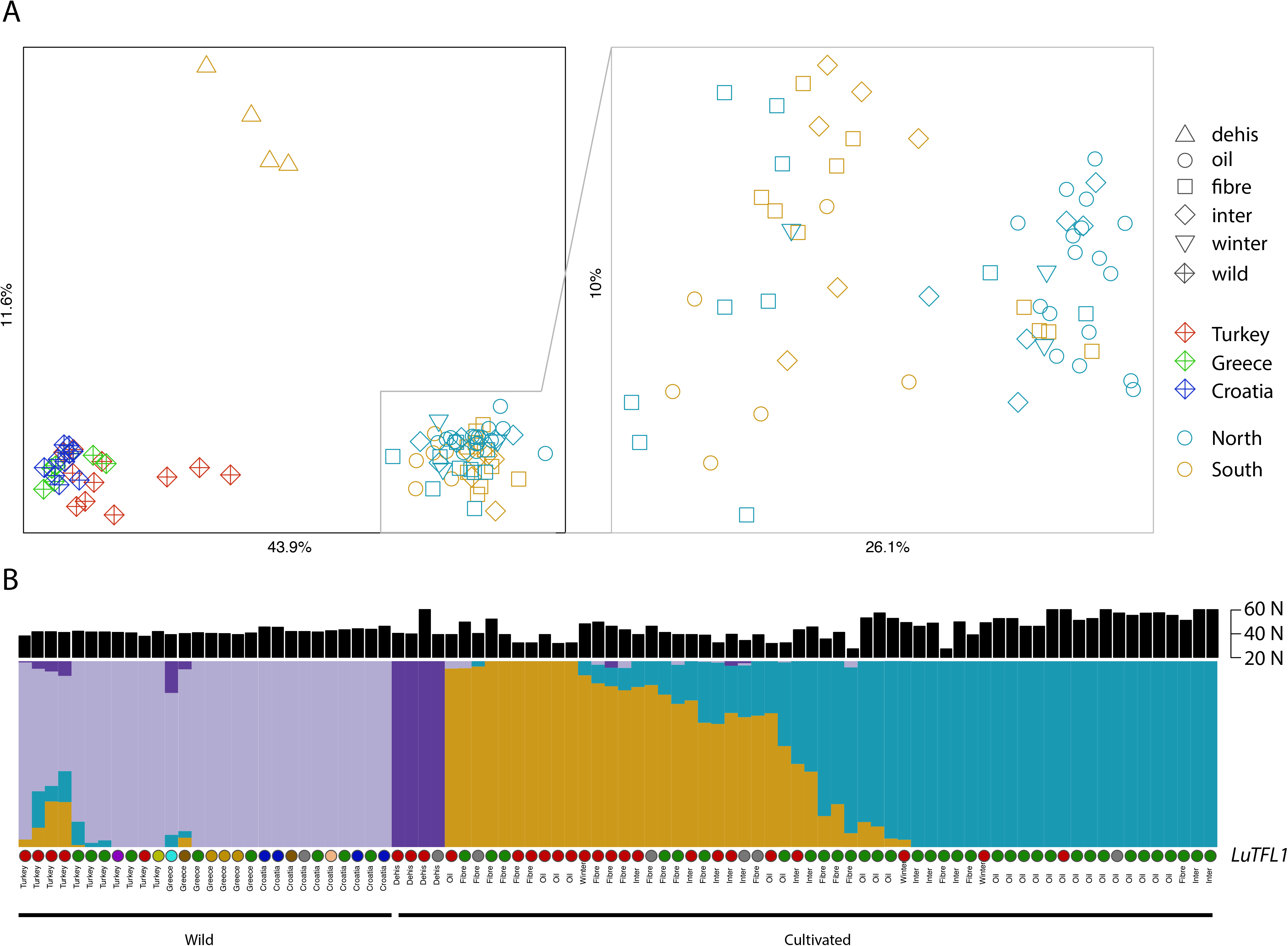
Population structure of pale flax and varieties of cultivated flax based on RADseq data: A. Multidimensional Scaling Plot of pairwise nucleotide distances between flax accessions separates three main flax populations. B. Admixture plot of flax four ancestral components. Pale flax samples are arranged from east (Eastern Anatolia) to northwest (Central Europe), while samples of cultivated flax were arranged based on admixture proportions. Above presented is the latitude at which the individual plant was sampled. Below, individuals are annotated with allele of *LuTFL1* they carry; colours are the same as in Figure 1.

Randomly sampled RADtags across the genome are a good proxy for neutrally evolving parts of the genome. We used allele frequencies of RAD SNPs in southern and northern flax populations as a null distribution for changes over latitudes due to population structure and demographic processes. We then tested allele frequencies of three *LuTFL1* SNPs, which distinguish allele I and III, against RAD distribution. Firstly, we compared fixation index statistics, Figure 3A. We found that the F_ST_ values for three SNPs associated with the differences between *LuTFL1* alleles I and III are significantly higher than the background range of values based on RAD loci (q-value 0.045). Secondly, we placed the position of the *LuTFL1* alleles on the two dimensional allele frequency spectrum of RAD loci (frequencies of northern population on x-axis; southern population on y-axis), and considered the relative deviation of allele frequency from the diagonal distribution, Figure 3B. We found that the three SNPs defining allele III lie significantly outside the background range (Supplementary Table S6), but close to the edges of the null distribution, whereas SNPs associated with allele XII are within the general range. Together, these data suggest that allele III was selected for as cultivated flax moved northwards.

**Figure 3.**
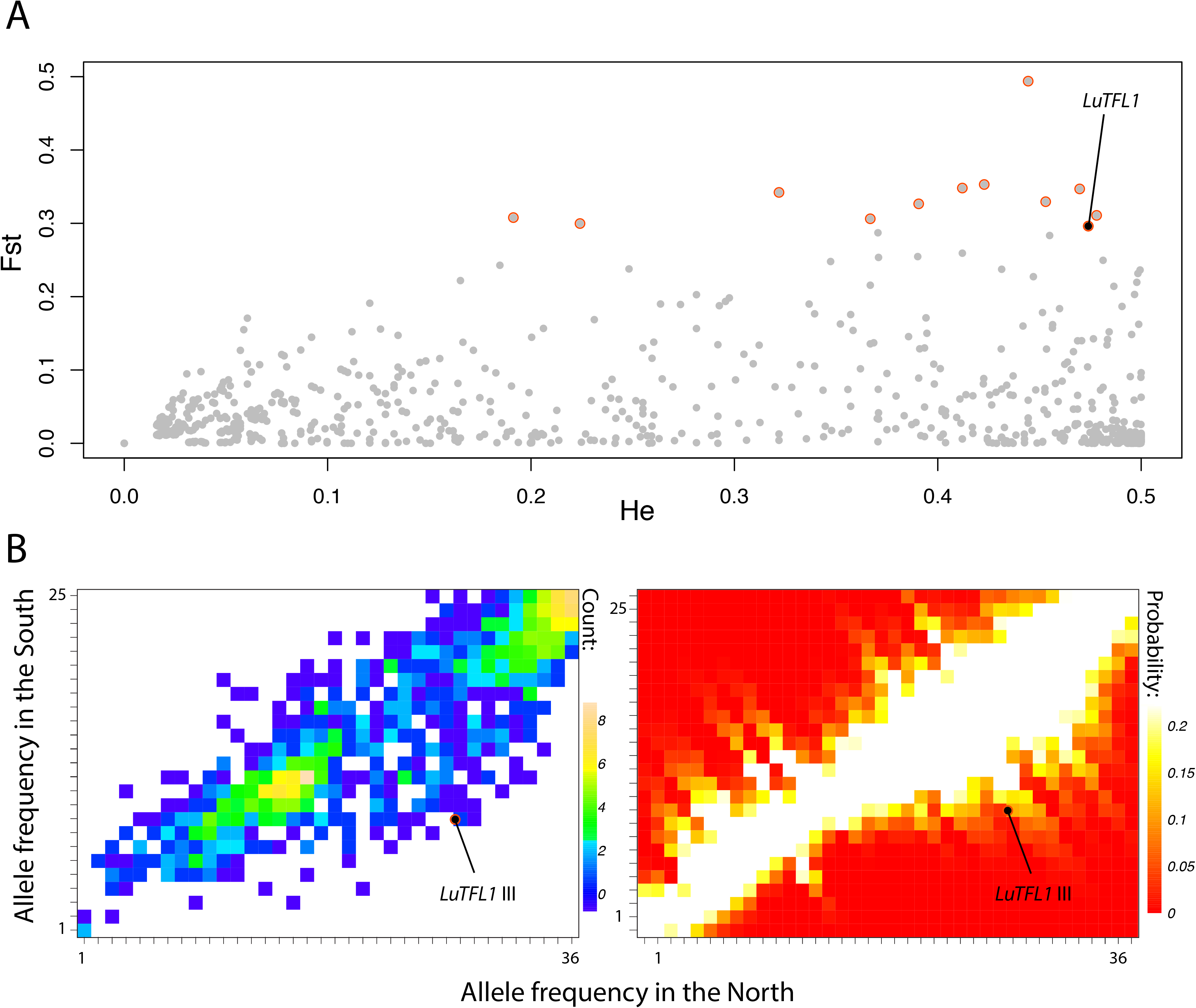
Outliers to neutral allele frequencies between northern and southern cultivated flax populations: A. F_ST_ values for RAD neutral SNPs plotted as a function of heterozygosity. Outliers with elevated F_ST_ and heterozygosity are marked with red circles. Outlier SNPs distinguishing alleles I and III in *LuTFL1* locus are marked with black. B. Two dimensional allele frequency spectrum of RAD loci alleles between northern and southern cultivated flaxes. Frequency of *LuTFL1*.III SNPs are marked as circles. To the right: probability heat maps of sampling frequency combinations given the allele frequency spectrum based on distance from the diagonal.

Given the proximity of the general differentiation of allele III between northern and southern populations relative to the null distribution evident in the allele frequency spectrum, we speculated that weak selection was probably involved. To investigate this further we applied a method to determine the most likely selection coefficient associated with allele frequency changes over time^39^, using archaeological dates of the arrival of agriculture at different latitudes to date the latitudinal allele frequencies, Figure 4. Our estimates confirmed weak selection with values of s generally between 0.001 and 0.007 (mean 0.003), but interestingly we found an increase in selection strength over time and latitude, indicating that allele III was progressively more strongly selected for as flax moved to higher latitudes.

**Figure 4.**
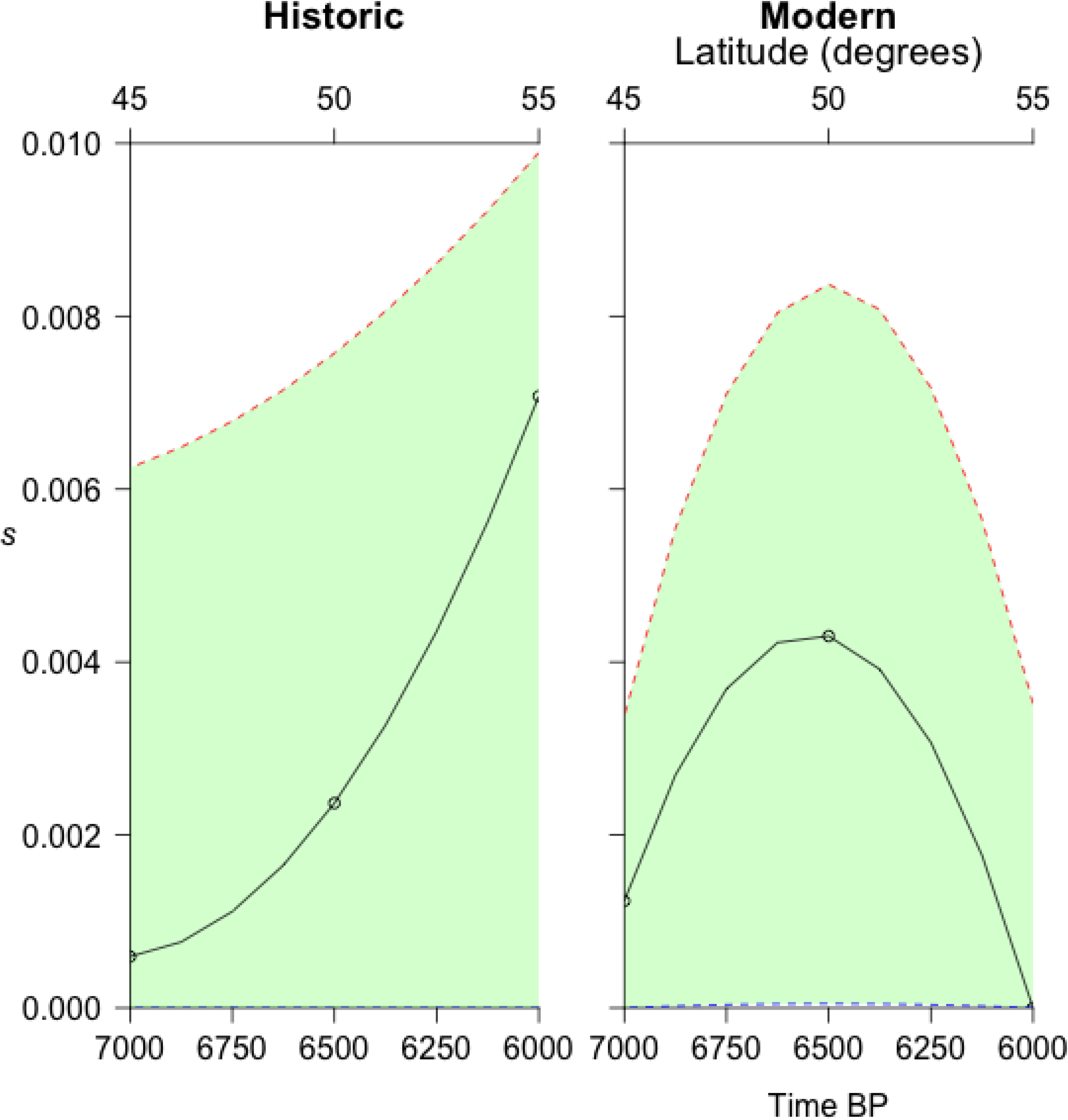
Strength of selection on *LuTFL1* III in modern and historic flaxes. Estimates of selection coefficient over latitude and time, where latitude correlates to time of arrival of agriculture.

Finally, we investigated whether the RAD loci data harboured evidence of movement of alleles between wild and cultivated flax populations beyond the area of domestication, which we implied from geographic distribution of *LuTFL1* alleles in wild and cultivated flaxes (Figure 1B, 1C). We observed no clear signal using f_3_ statistics (cultivatedN; cultivatedS, wildN), however, varying rates of population differentiation through founder effects and multiple complex processes of gene flow can mask signals of gene movement between populations^40^. We therefore considered portions of the data where evidence of gene movement may be clearer. We identified subsets of alleles that were discriminatingly higher at a range of frequencies in either northern or southern populations of wild flax. We reasoned that if cultivated flax had spread northwards without contact with wild populations, then the differentiation between northern and southern populations at these subsets of loci should not be perturbed relative to the background distribution of loci differentiation. We found that there was no significant difference between the cultivated populations at loci that were higher in southern wild populations, but that there was an elevated level of differentiation between cultivated populations at loci that were differentially high in northern wild populations (Supplementary Figure S7). These findings are verified by an increased ancestral information content (*I_a_*)^41^ at loci that were differentially high in northern but not southern wild populations (Supplementary Table S7). This indicates a bias against the neutral expectations of uncorrelated drift between the cultivated and wild population pairs that could represent gene flow from northern wild populations into northern cultivated populations. Alternatively, this signature could be an effect of a parallel selection for traits associated with high frequency RAD loci alleles in the wild population. We deemed the latter unlikely since 78% of loci that had elevated frequencies in northern wild populations also had elevated frequencies in northern cultivated populations, which would require an explanation of selection being responsible more frequently than drift for high frequency alleles in the north.

### Functional homology of *LuTFL1*

We explored the function of *LuTFL1* and *LuTFL2*, first by investigating expression of these genes in the leaf and shoot apex tissue of pale flax plants from six populations over the course of the plant development to flowering using a semi-quantitative PCR approach. *LuTFL1* was expressed continuously from the first point of measurement (40 days) to flowering (Supplementary Figure S8) consistent with *TFL1* expression in *A. thaliana^42^* as opposed to *ATC*, which is expressed in the hypocotyl in *A. thaliana^43^*. We detected no expression of *LuTFL2* in shoot tissue. Based on phylogenetic evidence, *LuTFL1* is more similar to *ATC*, however characterization of its expression pattern led us to conclude that *LuTFL1* is most likely functionally orthologous to *TFL1*.

We tested the hypothesis that *LuTFL1* functions as *TFL1* by complementing *TFL1* knock-out mutants of *A. thaliana* with a *LuTFL1* construct under a 35S promoter. We obtained a stable line after transforming a *tfl1-2* mutant in Landsberg *erecta* (Ler) genomic background^44^ using an *Agrobacterium* system. Over 160 plants of 35S::*LuTFL1* inbred progeny (T2) were grown in long day conditions. For all the plants we measured days to bolting, days to open flowers, days to end of flowering and stem height at the end of flowering. Flowering time phenotypes segregated in T2 with a Mendelian 15 to 1 ratio (Supplementary Figure S9), which suggests that two copies of transgene were integrated in *tfl1-2* mutant. Transgene presence/absence was validated using primers specific to the transgene in 21 plants with extreme phenotypes, Supplementary Figure S9. Plants that contained the 35S::*LuTFL1* transgene were compared with *tfl1-2* mutants and Ler-0 wild type. The start and end of flowering was significantly lower in *tfl1-2* mutants than in Ler-0 and 35S::*LuTFL1* plants (Figure 5; Supplementary Figure S9), suggesting that the flax *LuTFL1* I allele can rescue non-functional *TFL1* in *A. thaliana*. Constitutive expression of this allele under 35S promoter further delays flowering, when compared to Ler-0. Subsequently, the amount of expressed *TFL1*/*LuTFL1* has a positive impact on plant height at the end of flowering. We conclude that, similar to *TFL1* in *A. thaliana*, *LuTFL1* functions in delaying flowering, promoting indeterminate growth and finally resulting with increased plant height at senescence. The experimental data from transgenic 35S::*LuTFL1 A. thaliana* plants support the notion that the correlation between allele type and phenotype could have a causative relationship.

**Figure 5.**
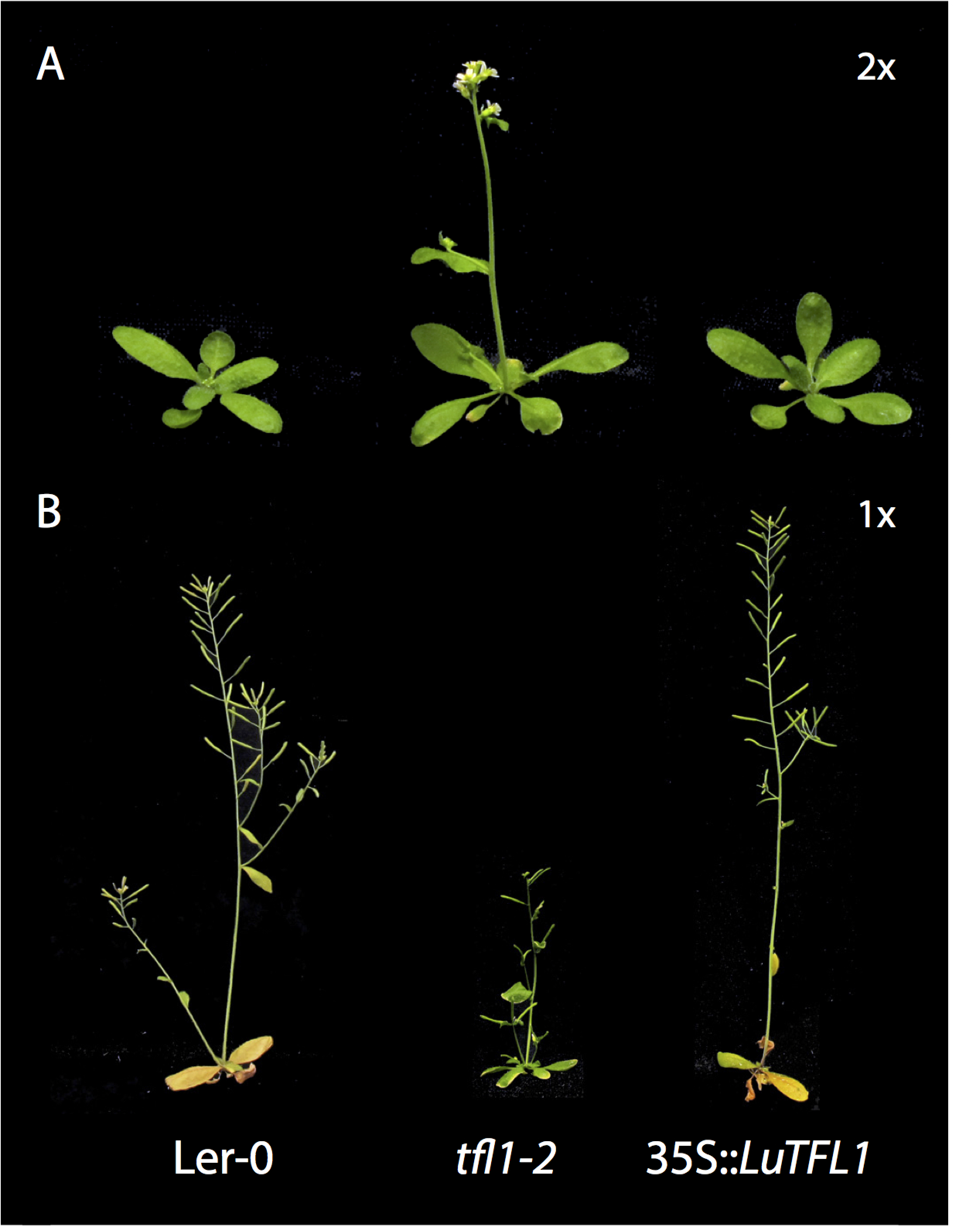
Flax *LuTFL1* gene complements *TFL1* mutant in *Arabidopsis thaliana*: A. Images of plants in 2x magnification at the median time to first open flower in *tfl1-2* mutant (20 days). Both Ler-0 wild type and tfl1-2 mutant complemented with 35S::*LuTFL1* are not yet flowering at that time. B. Plants at their median time to end of flowering in respective cohorts. Mutant *tfl1-2* ended flowering after 23 days and reached average height of 69 mm; by contrast Ler-0 wild type and 35S::*LuTFL1* plants flowered until 36th and 40th day reaching 134 and 211 mm height respectively.

Although *LuTFL1* is a functional homolog of *TFL1*, we still do not fully understand the effect of allelic diversity in flax at this locus. We detected no non-synonymous substitutions between groups I and III that could account for a potential functional basis for the difference in selection histories between the alleles. However, a transcription factor-binding site was identified to be present in the promoter regions of *LuTFL1* III and VIII, but absent from *LuTFL1* I (Supplementary Figure S4B). We hypothesized that such a difference between the allele groups could result in a change of expression pattern of *LuTFL1* and in consequence, altered flowering time, which in turn could be under selection in the northerly latitudes. To help understand how an adaptation to latitude through the *TFL/FT* system could produce the observed flowering and architectural changes in flax we applied a plant development model. The model was used to explore how selection at the *TFL1* locus and latitudinal movement could affect stem height, which correlates with increased fiber content^45^. The model predicts that increased *TFL1* expression is expected to produce longer stemmed plants, but also that at higher latitudes such plants would be more fit through a greater rate of flower production relative to long stemmed forms in the south (Figure 6). This model outcome predicts that adaptation to variable climate in northerly latitudes by indeterminate flowering through selection at a *TFL1* locus allows increase in stem height, which is associated with an improvement in fiber content.

**Figure 6.**
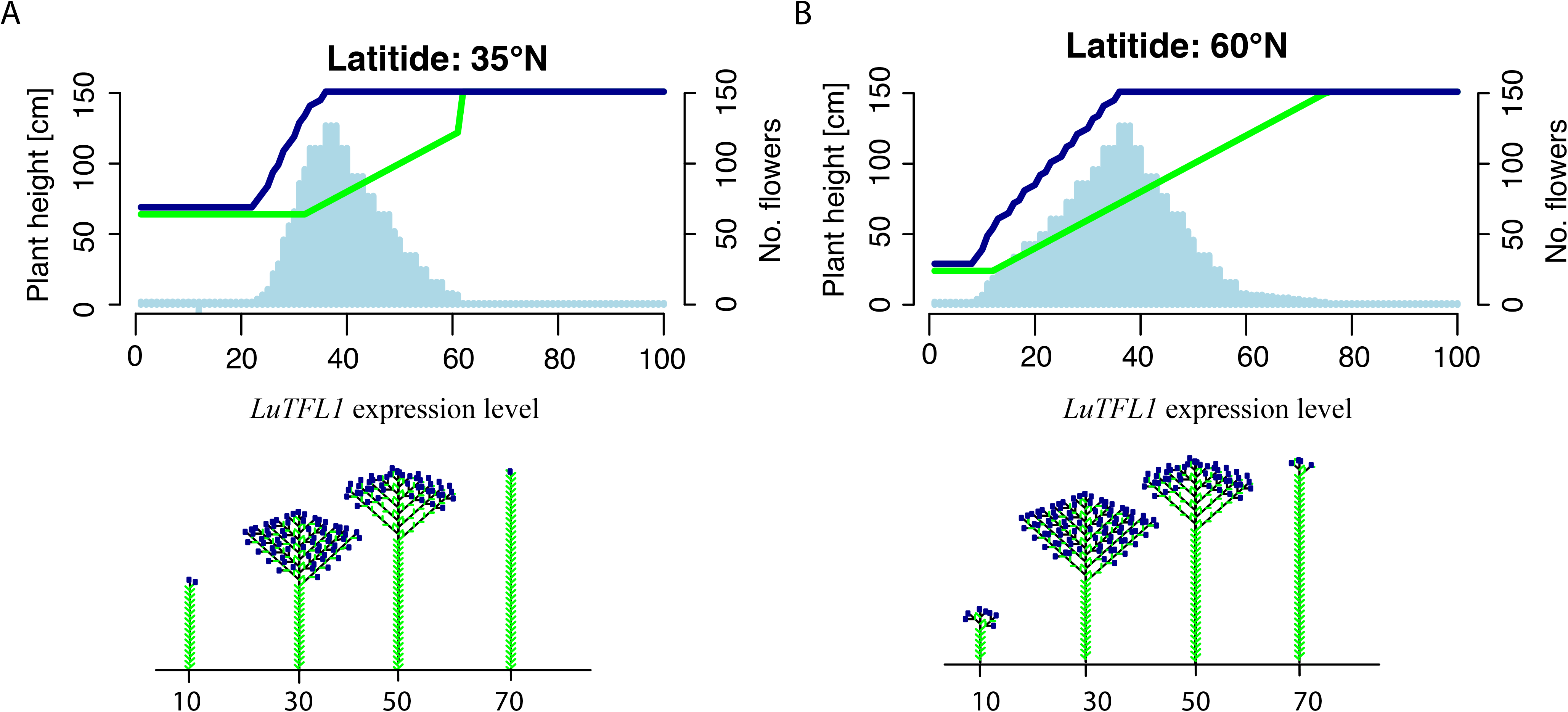
Plots summarizing parameters of flax inflorescence simulated with pgrowth model over different base levels of *TFL1* expression. Shown below charts are representative flax architectures associated with base levels of *TFL1*. A. Chart and architecture diagrams for plants grown at N 60° latitude with assumed floral initiation threshold set to 14 hours. B. Chart and architecture diagrams for plants grown at N 35° latitude with assumed flowering threshold set to 14 hours.

## Discussion

Crop plants were domesticated independently around the world at the ‘Domestication Centers’ in prehistoric times^1^. Early farmers spread out from those centers and brought with them major crops as they expanded into new geographic zones. There are numerous examples of crop plants that adapted to new environments through changes in flowering time, particularly during northwards expansions. Maize, domesticated in Mesoamerica, would only be successfully cultivated in the temperate North America after adapting through reduction of flowering time^46^. Similarly, after domestication of rice in Eastern China, its successful cultivation in Korea and Japan required changes in flowering behaviour^47^. In many plants, *TFL1* family genes that influence flowering time by acting antagonistically with *FT* to delay flowering^48–50^ have been identified as targets for selection in adaptation to northern latitudes and have also been shown to influence plant architecture^26,30,31^ and fruit yield^29^. *TFL1* homologs have played an important role in adaptation to temperate climate in barley^10^ and soybean^51^, where it has been noted that adaptive variant at this locus is strongly enriched in the northern end of crop distribution.

Cultivated flax was domesticated in the Near East alongside wheat and barley^2^ and likewise adapted to temperate climate of prehistoric Europe. In this paper, we show that flax genome encodes multiple PEBP orthologs. Of those, we show that *LuTFL1* has latitudinally structured allelic diversity and molecular signatures consistent with selection. This has been validated against null distribution of neutrally evolving loci, which we generated using RADseq approach. To further reinforce evidence of selection on *LuTFL1*.III in the northern Europe, we anticipate the availability of full genome information for flax populations. Similarly to other crop plants, adaptation to Northerly latitudes in flax was achieved through modifications in flowering behaviour. Through heterologous overexpression of *LuTFL1*, we show that this gene is in fact a functional homolog of *TFL1* and is capable of changing flowering time^26,30,52^. Similarly, *LuTFL1* is shown to be a decisive factor in establishing plant architecture and height. Based on that we conclude that the increase in frequency of *LuTFL1*.III in the north and signatures consistent with natural selection are results of flax adaptation in Europe through changes in flowering time. Interestingly, this change had an impact on flax architecture and could have predisposed flax to fibre production. This evolutionary transition may have been key to the rise of the textile revolution in central Europe^20^, which is evidenced in the reduction of flax seed size^19^ and improved tools for fiber production^21^ in Neolithic contexts. Our conclusions are consistent with phenotypic associations in cultivated flax, in which fibre content is correlated with plant height and flowering time at the same time^45^.

Flax is shown to have adapted to Northerly latitudes in a similar fashion to other crop plants. However, this study additionally provides an unusual example on how the influence of natural adaptations within a crop may ultimately influence the use of that crop. Normally, the evolution of domesticated plants is associated with an initial rise of a suite of traits collectively termed the domestication syndrome^53^, later followed by diversification or improvement traits^54^. Such traits are considered for the most part to be agronomically advantageous. Mounting evidence shows that plant adaptations within the human environment are not restricted to crops^55–57^ and within crop plants not all adaptations are of obvious benefit to agronomic productivity and yield, but related to the wider ecology^58^. Examples include a tendency to increase rates of water loss in domesticate forms^59^. Such adaptations highlight a tension between the effective exploitation of plants as a resource by endowment of useful traits and ongoing natural adaptations that occur in plants that may compromise productivity. This study shows that the adaptation of flax to the European variable climate during Neolithic likely involved the *LuFTL1* locus and resulted in a change of cultivation purpose through substantial modification of the stem height, which is correlated with the technical fiber quality.

Natural adaptations in crops are commonly acquired from wild relatives’ standing variation. One case of such transfer is the Mexicana teosinte introgression into highland maize landraces^60^. It is notable that flax natural adaptation appears to have occurred through a transmission from the wild to the cultivated gene pool despite the highly selfing nature of both cultivated and pale flax, at only 3-5% outcrossing. Furthermore, wild flax stands are typically of low density. It is possible in this case that entomophilous pollination could have played an important role either by natural pollinators or even the action of domesticated bees of the early farmers^61^. Further research is required to understand the interplay of these evolutionary forces and how they contribute to the tensions between artificial and natural selection.

## Materials and Methods

### Plant Materials

Samples from 16 populations were collected from natural habitats in the summer of 2011 from Croatia, Montenegro, Albania, Greece and Bulgaria. These were combined with a further 16 populations of pale flax supplied by the Plant Genetic Resources of Canada, Agriculture and Agri-Food Canada, Saskatoon Research Centre. Seed material for 58 modern cultivars of flax and 18 landraces were obtained from the Plant Genetic Resources of Canada, Agriculture and Agri-Food Canada, Saskatoon Research Centre. A further 28 historic landrace accessions were obtained from Plant Genetic Resources of the N. I. Vavilov Research Institute of Plant Industry, St. Petersburg, Russia. Samples ranging in age up to 194 years were sampled from Herbaria variously from the Natural History Museum, University of Oxford and the University of Warsaw.

### Identification and resequencing of *LuTFL1* locus

Based on multiple alignments that contain sequences from different eudicots, degenerate primers were designed in exonic regions to cover between 400 bp of *TFL1* region. The putative sequences of *TFL1* in flax were subject to BLAST searches against the cultivated flax genome scaffolds v1.0 and CDs v1.0 databases^62^ in order to identify full sequences of genes of interest and their flanking regions. Phylogenetic trees with *TFL1* homologs from *A. thaliana* and *P. nigra* were estimated using Bayesian Inference in MRBAYES v3.2^63^.

A total of 148 samples were used in this study, including 58 cultivars, 18 landraces, 38 historic landraces of cultivated flax, 32 pale flax samples and two *L. decumbens* accessions. The total DNA from 20 mg of seeds was isolated using modified DNEasy^®^ Plant Mini Kit (Qiagen) protocol. In case of herbarium specimens – mixed plant tissue was used for extraction using CTAB protocol^64^. A 1367 bp target region was amplified from *LuTFL1* spanning exons 1-4 using primers (5-TTACAACTCCACCAAGCAAGTC, and 5-TGTCTCGCGCTGAGCATT). Haplotype networks was made using uncorrected_P character transformation and Rooted Equal Angle splits transformation in *SplitsTree4^65^*. The size of the network nodes was made proportional to the number of samples that share a particular haplotype. The relationship between latitude and *LuTFL1* and *LuTFL2* haplotypes separately was modelled using logistic regression in *R* programme and plotted using *popbio* package.

### Selection signatures at *LuTFL1* locus

In order to test if *LuTFL1* and *LuTFL2* were inherited neutrally or were under selection, neutrality tests with variety of estimators were carried out. Tajima’s D^66^, Fu and Li’s D, Fu and Li’s F^67^ statistics were calculated in *INTRAPOP* and the R2 statistic^68^ was calculated in *R* using the *PEGAS* package. The extent of linkage disequilibrium was examined between the alleles of the *LuTFL1* and *LuTFL2* loci. Expected frequencies of genotypes were calculated from the observed individual allele frequencies. The correlation coefficient of *D*, *r*, was calculated from observed genotyping data where:

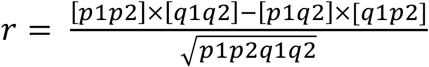

Square brackets indicate the observed proportion of the sample of genotype combinations, *p1, p2, q1 and q2* refer to the allele frequencies of *p* and *q* type alleles at the first and second locus respectively, and the numerator equates to the statistic *D*.

Each latitude in Europe is associated with a different time of arrival of agriculture based on archaeological data. The program SELECTION_TIME.pl^39^ estimates selection coefficients from dated frequencies. The model takes as input a series of dated frequencies, mating strategy of the organism, whether the data is phenotype or allele frequency and whether the trait under selection is dominant or recessive. The program takes a pair of input observed dated allele frequencies as start and stop frequencies. The inbreeding coefficient *F* is used to determine the initial genotype frequencies. The selection coefficient *s* is then described by equation:

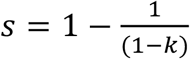

Where *k* is the selection differential. To account for sample error, we generated upper and lower bounds of *s* with Beta distributions assuming a binomial process of *x* observations in *n* samples, where x equates to the number of samples carrying the *LuTFL1* III allele, and *n* equates to the total number of samples at a specific latitude. For details see Supplementary Methods.

### RAD sequencing and processing

From the total collection, 90 plants were chosen for the RADseq experiment (Supplementary Table S1). They represent 28 pale flax and 62 cultivated flax accessions. Plants were grown in glasshouse conditions and harvested after seedlings reached 10 cm of height. Snap-freezed material was ground with 3 glass beads (3 mm diameter) in TissueLyser machine (Qiagen) and then used for DNA isolation with DNeasy^®^ Plant Mini Kit (Qiagen) following the manufacturer’s manual. DNA was digested with 2U of SbfI HF restriction enzyme (New England Biolabs) for one hour and then the reaction was heat inactivated. For RADseq library preparation, 40 µl of genomic DNA at concentration of 25 ng/µl was prepared following previous methods^34^. Barcoded libraries of 10 different DNA isolates were pooled together into the total of 2 µg of DNA in 300 µl of solution. DNA was sheared ten times on ice in Bandelin Sonoplus HD 2070 sonicator. Sheared DNA was purified using AMPure XP beads (Agencourt) in proportion 1 to 1 with buffer. Approximately 20 ng of library was amplified with Phusion HF Master Mix (NEB) in total of 14 cycles. The first 50 libraries were sequenced on HiSeq 2000 Illumina platform at Oxford Genomics Centre using the TruSeq reagents. Another 40 samples were submitted for sequencing on Genome Analyzer II Illumina platform in Genomic Centre at University of Warwick.

Raw reads from Illumina sequence runs were assembled into FastQ format and assessed in FAST QC software v0.10.1 to check for standard quality measures: quality scores, overrepresented sequences, GC content and N content^69^. We generated SNPs from *de novo* assembled RADtags. Scripts from the STACKS pipeline v1.05 was employed to de-multiplex and discover RAD sequence markers^35^. Low quality sequences and those characterized by erroneous barcodes were discarded. Remaining sequences were sorted according to their barcodes. SNPs were called with the following settings: 5 identical, raw reads required to create marker stack, 5 mismatches were allowed between alleles in single individual and calling haplotypes from secondary reads was disabled.

### Population structure and differentiation

Distances calculated from identity by sequence between all individuals were measured using plink (v1.07)^70^. This distance matrix was used for multi-dimensional scaling (MDS) approach in R with cmdscale function. Population ancestry of each individual was modelled using ADMIXTURE (v1.23)^36^. We assumed existence of two to seven ancestral populations and chose the model, which was characterised with the lowest five-fold cross-validation error.

We utilized the ancestral information content concept^41^ to employ a high variant test to explore possible gene movement between populations when there may be complex confounding movements masking a general signal. We identified three subsets of loci in which the difference in frequency (∂f) between northern and southern wild populations was more than 0.5, 0.4 or 0.3, with the higher allele in the northern population. We examined the ancestry information content of the ∂f loci subsets using the *I_a_* (*I_a∂f_*) statistic relative to the information content of all loci (*I_at_*). We then compared the ∂f subset mean F_ST_ with the total null background of markers 100000 randomly sampled subsets, for details see Supplementary Methods.

Cultivated flax individuals were segregated into two groups based on their latitude; individuals from above N40° were included in the northern subpopulation while individuals from below N40° in southern. We calculated size-corrected and uncorrected F_ST_ values between northern and southern cultivated flax populations for each RADtag using functions in OutFLANK R package. We applied trimming approach to infer the distribution of F_ST_ for neutral markers and used likelihood estimation for outliers^71^. Allele frequency spectrums were generated from in house scripts from STRUCTURE format files acquired from *de novo* assembly RAD tags. P value heat maps were generated from the allele frequency spectrums using in house scripts that calculated probability density functions of Gaussian distributions from mean and standard deviations of frequency distributions perpendicular to the diagonal.

### Expression and functional homology of *LuTFL1*

Plants of six populations (W042, W043, W067, W069, W077 and W094) were sown in 100 replicates each. Seeds were stratified for 3 days in the dark at 4°C and grown at room temperature with a 16-hour photoperiod for 10 days followed by vernalization in cold room at 4°C for a period of 40 days. Subsequently plant were moved to growth chambers with 16h daylight at 24°C until they flowered. Samples of three plants were taken at 0, 15, 17, 19 and 21 days. Samples were snap-frozen and the total RNA extraction was carried out from 20 mg of ground tissue using mirVana™ miRNA isolation kit (Invitrogen) following the standard protocol. DNA contaminants were digested in reaction with 2 units of DNase I (Invitrogen) and 2 µg of total RNA in DEPC-treated water. cDNA was synthesized from 1µg of DNase-treated RNA with use of SuperScript^®^ II reverse transcriptase and unspecific Oligo dT primer following the manufacturer instructions. Specific primers were designed in exonic regions of *LuTFL1*, *LuTFL2* and *LuFT* to cover approximately 150 bp long fragment. The *GAPDH* housekeeping gene was chosen as an expression control. Semi-quantitative PCR was carried out for each sample with 30 cycles.

We have used Landsberg erecta (Ler-0) ecotype with functional *TFL1* gene as a reference and its tfl mutant, *tfl1-2^44^* for complementation experiment with *LuTFL1*.III coding sequence. In these experiments plants of *Arabidopsis thaliana* were grown on soil in long days (16 h light/8 hours dark) under a mixture of cool and warm white fluorescent light at 23°C and 65% humidity.

Coding sequence of *LuTFL1*.III allele was amplified and introduced into a Sma1-digested Gateway entry vector, pJLBlue[rev], as a blunt-end fragment. The resulting entry vector, propagated in *Escherichia coli* DH5α strain, was then used in a recombination reaction with a modified Gateway recombination cassette pGREEN-IIS^72^. This recombination reaction effectively placed the *LuTFL1*.III allele behind the constitutive CaMV 35S promoter in the pGREEN vector conferring resistance to BASTA. Sequence of PCR products and subsequent plasmid vectors were checked by Sanger sequencing and compared to the expected input sequence from the cDNA and vector backbone. The expression construct was introduced into *A. thaliana tfl1-2* mutants by *Agrobacterium tumefaciens*-mediated transformation^73^. Transformed seeds were stratified and selected in soil with BASTA herbicide. Seeds from a single line with resistance to BASTA were collected.

Second generation of 35S::*LuTFL1*.III tfl1-2 transformants was sown together with wild type Ler-0 and *tfl1-2* mutant plants for phenotyping. We have measured flowering time as days to emergence of floral meristem and days to opening of first flower. Additionally, we measured days to end of flowering and plant height at the end of flowering. We have selected random 32 individuals for transgene genotyping. The difference in mean phenotypes for 35S::*LuTFL1*.III, Ler-0 and tfl1-2 was tested using t-test in R.

Growth models under different *LuTFL1* expression levels simulated using in-house R script pgrowth, for details see Supplementary Methods.

## Acknowledgments

We would like to acknowledge Sabine Karg, for her help in clearing the archaeological background for this study and help in obtaining historic samples from Vavilov Institute, Sankt Petersburg. We would like to thank Nina Brutch (Vavilov Institute) and Dallas Kessler (PGRC) for dispatching seeds from seed banks. Toni Nikolic and Arne Strid helped in locating pale flax populations in the Balkans. Mariusz Czarnocki-Cieciura, Agnieszka Cakala and Ewa Samorzewska helped during pale flax collection. Finally, we would like to thank Hernan Burbano and Detlef Weigel for support during revisions and functional analyses. RMG was supported by University of Warwick Chancellor’s Scholarship scheme, OS is supported by NERC (NE/L006847/1) and RW is supported by NERC (NE/F000391/1). Sequence data has been entered into GenBank: *LuTFL1* (KU240116 - KU240259) *LuTFL2* (KU240260 - KU240389), RADseq (PRJNA304385).

## Author contributions

RMG, MZ, OS and RW carried out the research. RMG, YBF and RGA designed the project. RMG and RGA analyzed the data. RMG, YBF, AD and RGA wrote the manuscript.

